# Individualized discovery of rare cancer drivers in global network context

**DOI:** 10.1101/2021.10.04.463007

**Authors:** Iurii Petrov, Andrey Alexeyenko

## Abstract

Late advances in genome sequencing expanded the space of known cancer driver genes several-fold. However, most of this surge was based on computational analysis of somatic mutation frequencies and/or their impact on the protein function. On the contrary, experimental research necessarily accounted for functional context of mutations interacting with other genes and conferring cancer phenotypes. Eventually, just such results become “hard currency” of cancer biology. The new method, NEAdriver employs knowledge accumulated thus far in the form of gene interaction networks and functionally annotated pathways in order to recover known and predict novel driver genes. The driver discovery was individualized by accounting for mutations’ co-occurrence in tumour genomes. For each somatic genome change, probabilistic estimates from two lanes of network analysis were combined into joint likelihoods of being a driver. Thus, ability to detect previously unnoticed candidate driver events emerged from combining individual genomic context with network perspective. The procedure was applied to ten largest cancer cohorts followed by evaluating error rates against previous cancer gene sets. The discovered driver combinations were shown to be informative on cancer outcome. We demonstrate that the individualized discovery revealed driver events which were individually rare, not detectable by other computational approaches, and related to cancer biology domains poorly covered by previous analyses. Considering the novel driver candidates and their constellations in individual tumor genomes opens a novel avenue for personalized cancer medicine.

## Introduction

Carcinogenesis is a complex, multi-step process, during which cellular genomes accumulate somatic alterations. Mutations that cause or facilitate cancer initiation and progression are called drivers. On the other hand, many mutations occur spuriously due to impairment of chromosome maintenance, replication errors etc. Therefore, a cancer cell genome usually represents a mixture of driver and passenger mutations(1). Apart from the boosting genome instability that generates passengers as well as additional drivers, cancer cells should acquire selective advantages, such as apoptosis evasion, unconstrained proliferation, or survival in low-oxygen environment which correspond to the hallmarks of cancer(2). Given the avalanche of new data from cancer genome sequencing, it became possible to complement earlier known, “core” cancer gene sets with multitudes of computationally inferred drivers. Most of such novel sets are generalized either globally, within the pan-cancer paradigm(3,4) or within site/organ specific tumour types(5), sometimes delineating subtype-specific drivers(6),(7). This situation apparently contradicts the individualized approach to cancer treatment, which suggests molecular pathological analyses for disease prognostication, administration of targeted drugs(8), and discovery of novel drug targets. Currently, the majority of patients are not amenable to any approved targeted treatment since respective matching mutations occur with low prevalence. Further development of precision cancer medicine requires considering functional context of cancer genome in each patient(9). We pursue this approach via network analysis of mutated genes, by which patient-specific driver constellations shall be discerned from the background of passenger burden.

The existing computational approaches to cancer driver discovery can be classified into three major groups:

1. Mutation frequency analyses, based on the idea that driver genes appear mutated more often than expected by chance(10),(11). In order to keep discovering novel drivers, frequency methods should capture increasingly more rare events(12) – which is limited by practically achievable genomics dataset sizes. As an example, MuSiC driver analysis included only point mutations (PM) that occurred in more than 5% of tumours(13). A close-to-comprehensive frequency analysis might require 600-5,000 samples per tumour type, depending on background mutation frequency(14). Another problem is lower statistical power with regard to shorter genes. We found that short genes were underrepresented in all alternative sets considered in this study, except curated KEGG pathways. Meanwhile, even rarely mutated genes can be drivers e.g. in absence of alterations in a “major” gene, such as TP53 – but identifying such associations would require genome-wide studies at an unaffordable scale(15). Thus, despite all the advancements, cancer sequencing often fails to identify any driver events in a certain cancer genome.
2. Evaluation of functional impact of sequence alterations using protein structural information, physicochemical features, evolutionary conservation etc.(16),(17),(18) Such methods might also include frequency analyses and were often trained on smaller sets of best known cancer genes(19) which might lead to overfitting. Although some positive correlation with higher mutation frequency has been demonstrated (20), predictions by different methods often disagreed even for most studied genes(21).
3. Commonality of protein function to disease genes established via expert judgement or computational analysis of literature associations(22) and global gene network context(23),(24),(25). In contrast to the approaches described above, this “guilt-by-association” (GBA) methodology(26) did not require information on mutations *per se* and could thus be applied to all known genes. Most commonly, likelihood of a general function such as cancer “driverness” was assigned by a GBA algorithm alone which, when applied to all the genes, generated thousands of predictions with prohibitively high false positive rates (FPR).

A particular challenge would be to identify drivers among gene copy number alterations (CNA), which may encompass longer chromosomal regions with multiple genes were gained or lost at once, “competing” for a driver role assignment. Therefore, CNA genes were often excluded from the analyses described above.

Network analysis is an important tool for cancer driver gene discovery: it not only implements the GBA principle, but also assesses genomic events by employing the network-defined entities, such as modules and pathways. Non-biological algorithms of network analysis, such as PageRank(27) and Random Walk with Restart (RWR), exist since long ago and were adopted, usually with minimal or no changes, by bioinformatics frameworks(28–32). For a combination of natural and historical reasons, interpretation of these algorithms tend to be focused towards network hubs, which could miss novel disease genes with lower node degrees(33). Furthermore, GBA benchmarks might be incapable of finding truly novel genes, focusing on a well-known gene core, again mostly network hubs(34). This problem could be circumvented by ontology-independent methods, such as identifying modules of mutated genes that correlate by expression(23), physically interact(35), or share annotations(36). Such methods, though, when applied to either all or to frequently mutated genes would suffer from the high false positive rate or miss rare drivers, respectively. Another solution was offered by the method of network enrichment analysis (NEA)(37), where network connectivity is normalized by gene node degrees, which allowed studying genes poorly covered with experimental data. The other advantage of NEA is its high sensitivity and robustness due to considering the multitude of edges available in the global network(38),(39). The concept of enrichment, i.e. finding signal that prevails over noise in NEA is implemented via counting network edges that connect gene nodes. Significant excess of actual number of edges over what is expected by chance can distinguish functionally relevant genes, i.e. drivers differ from passengers by relevant fragments of network connectivity.

The above mentioned problem of high FPR can be efficiently addressed by combining probabilities from multiple evidence channels. In the presented analysis we did that in a three-pronged way. First, we reduced FPR by considering only genes altered in a given tumor genome, whereas genes mutated elsewhere were not evaluated. Second, we employed the idea that driver mutations of mutated gene sets (MGS) in individual samples should be mutually related and identified such cases by network enrichment against each other, i.e. within MGS. Third, we detected driver roles by summarizing network connectivity of MGS to a number of informative pathways. Since the full set of such pathways was not known in advance, we started from hundreds pathway profiles, followed by feature selection and creation of predictive cohort-specific sparse models. Combining the two predictors decreased FPR even further.

We applied the analysis pipeline to nine largest TCGA cohorts as well as to a newly compiled meta-cohort of medulloblastoma (MB). We evaluate agreement between our and earlier published driver sets, relative contributions of the driver score components, significance and error rates of predictions, and prove the method robustness across over a broad range of mutation rates (from very low in MB to very high in skin melanoma) and variable disease aggressiveness. Finally, we demonstrate functional relations between driver mutation patterns, gene expression in affected pathways, and patient survival.

## Results

### Algorithm outline: two evidence channels for driver prediction

The procedure evaluated likelihood of each genomic alteration (either PM or CNA) being a driver in each tumor genome. This was done by considering functional network context (Fig. 1) in two parallel, independent analysis channels:

**Figure 1.**
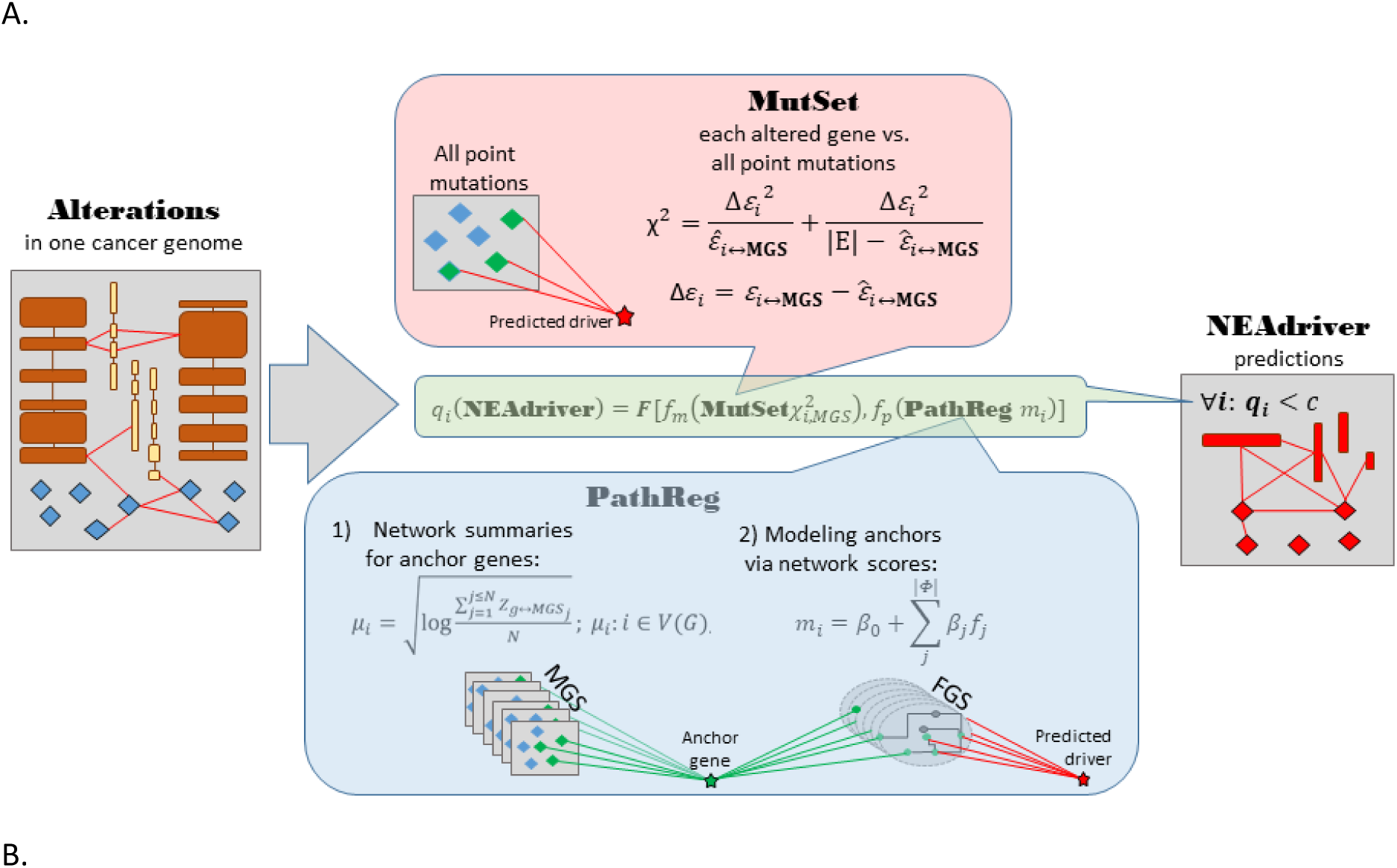
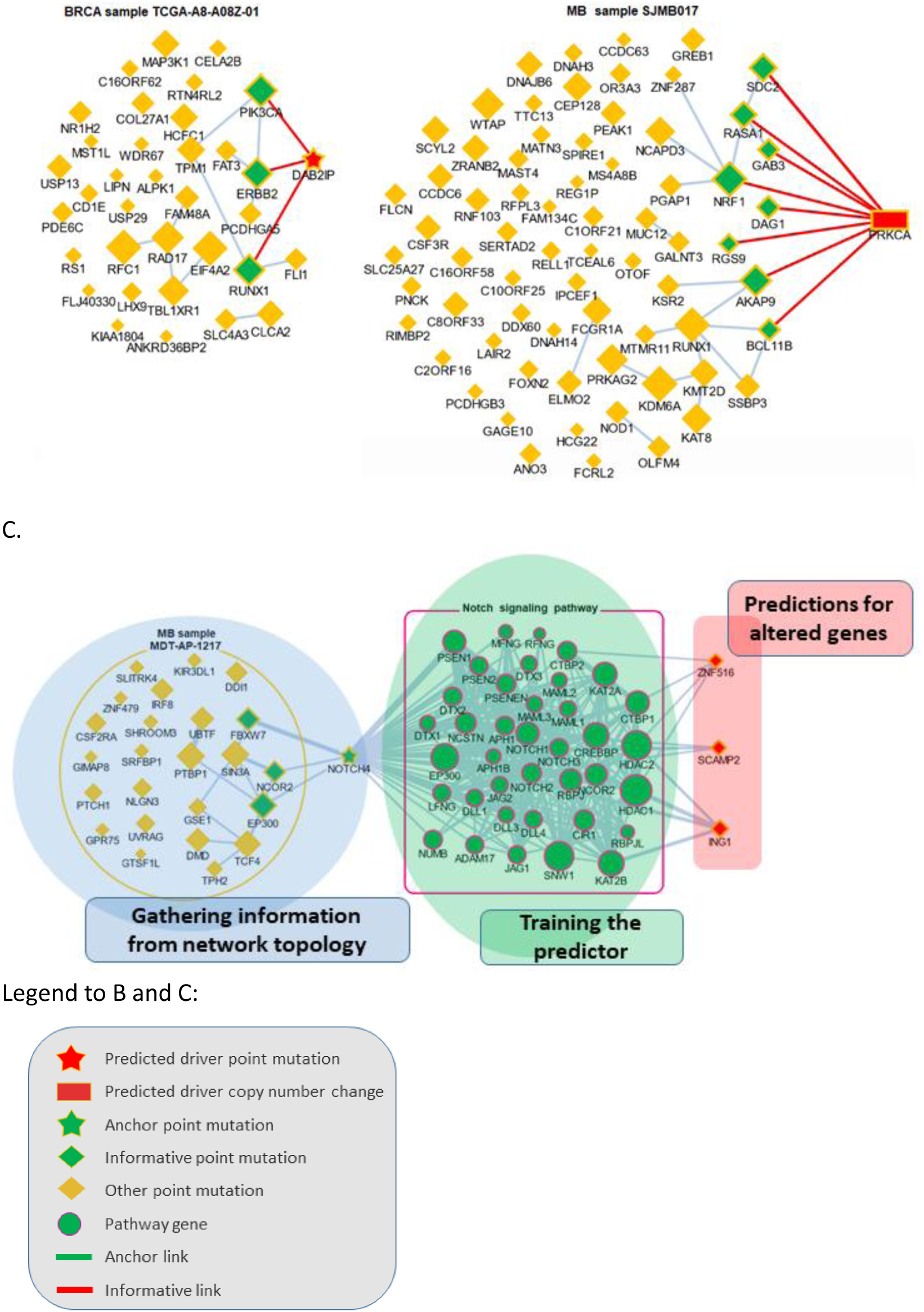
Visualization of NEAdriver analysis. A. Workflow according to the algorithm described in Methods. B. Examples of MutSet analysis to mutation gene sets in two cohorts. C. Example PathReg analysis in MB cohort. Legend to nodes and edges.

`MutSet` **channel** evaluated network enrichment between each altered gene *m* (*m* ∈MGS) and the constellation of all other altered genes *n* (*n* ∈MGS; *n* ≠ *m*). The resulting NEA scores *Z*_*m*↔*MGS*_ accounted for network degrees of the interacting MGS genes and expressed strength of the cumulative interaction compared to a value expected by chance, i.e. when *i* would be functionally unrelated to the rest of MGS.

`PathReg` **channel** evaluated likelihood of being a cancer driver in a two-step procedure. First, each of the *N* genes present in the network (*N*=19035) were characterized – in the same way as in MutSet – by network enrichment scores against each individual MGS. These “anchor” scores *Z*_*i*↔*MGS*_ were summarized per gene *i* and within each of the ten cohorts *c*. The reasoning behind this was that if MGSs included some actual drivers, then potential cancer involvement of a gene could be expressed as a summary of its network interactions over the MGS collection. For a highly scoring gene, this should become evidence of being a driver when it was altered in a given genomic sample. The cohort-specific gene vectors called `anchor.summary` should already contain all information gathered from genes’ interactions with all the MGSs. However, given that both MGSs and the available network were likely incomplete, some genes in `anchor.summary` would not be fully evaluated. Therefore, we introduced a second step: boosting via pathway enrichment scores. Cohort-specific predictive models were created using `anchor.summary` vectors as independent variables (being split into training and testing halves, *N’*=*N*/2 and N’’=*N*/2). Dependent variables for the models were provided by pre-calculated NEA scores for the same *N* genes against a collection of *P*=320 pathways, generally called functional gene sets (FGS), forming an *N*×*P* matrix. Then sparse multivariate models (k=31…83 pathways with non-zero coefficients) were obtained via the lasso training procedure under cross-validation and by controlling performance and robustness with error terms and information criteria. Due to big sample sizes (*N’* ∼ 10^4^), the models reproduced well on the test sets (Spearman rank R between actual and predicted `anchor.summary` vectors were 0.62…0.79; Supplementary Fig. 1). The sparse models produced cohort-specific `PathReg` scores for each of the *N* genes. Using published driver sets as references, performance of the scores was compared to the original `anchor.summary`, which demonstrated clear superiority of `PathReg` and thus gain in driver-related information via pathway enrichment. Thus, `MutSet` estimated driverness strictly in the context of individual cancer genomes, whereas `PathReg` models were originally derived from individual MGSs and then presented as universal, cohort-specific values. Importantly, neither of the two employed information on mutation frequencies. Genes that scored highest in `PathReg` are plotted as enriched against pathways included in the models (Supplementary Fig. 1).

The `MutSet` and `PathReg` scores were calibrated and converted into *p* and respective *q* (false discovery rate) values. Evidence from `MutSet` and `PathReg` was combined under OR condition, i.e. the resulting product q(M&P)=q(M)*q(P) reported the probability of NOT being a driver given evidence from the both channels. In this way, MGSs were reduced to driver gene sets, DGSs. For the purposes of further testing, the DGSs were defined at two significance thresholds q(M&P)<0.05 and q(M&P)<0.01. The full list of genes altered in the ten cohorts is presented in Supplementary Fig PathReg_and_combined_scores.pdf. We had also tested evaluation of mutations against sets of most frequently mutated genes (similarly to NetSig5000 method(40)), but this channel did not yield any advantage and was not used.

### Validation

We first evaluated performance of the method [q(M&P)<0.05] by ability to detect gene members of nine alternative reference sets, derived from either curated resources or computational analyses (Fig. 2A). Overlaps with the reference sets were mostly significant: 86 out of 90 pairwise comparisons by Fisher’s exact test and 58 out 90 by Mann-Whitney test received a Bonferroni-adjusted p-value below 0.05.

**Figure 2.**
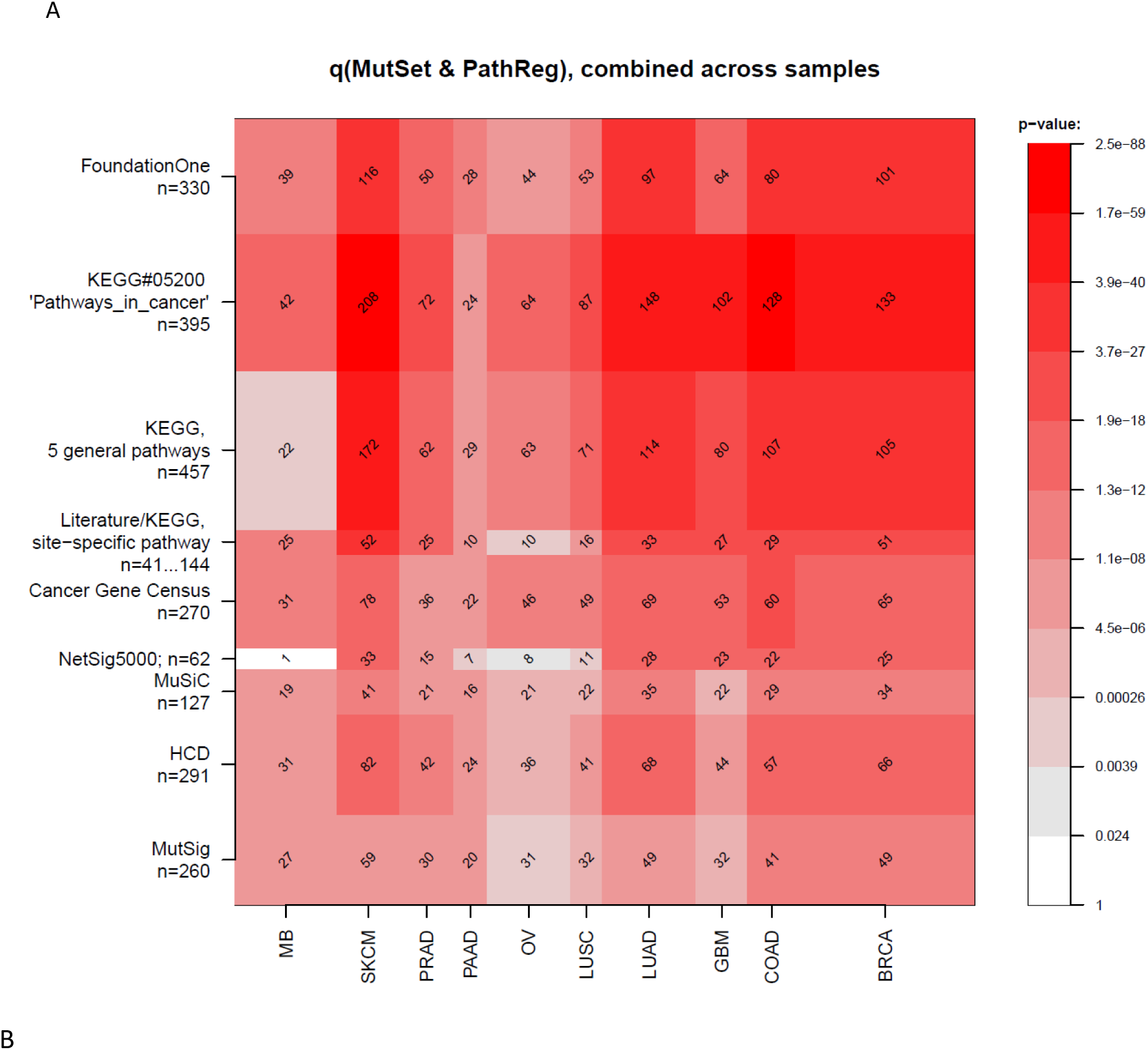
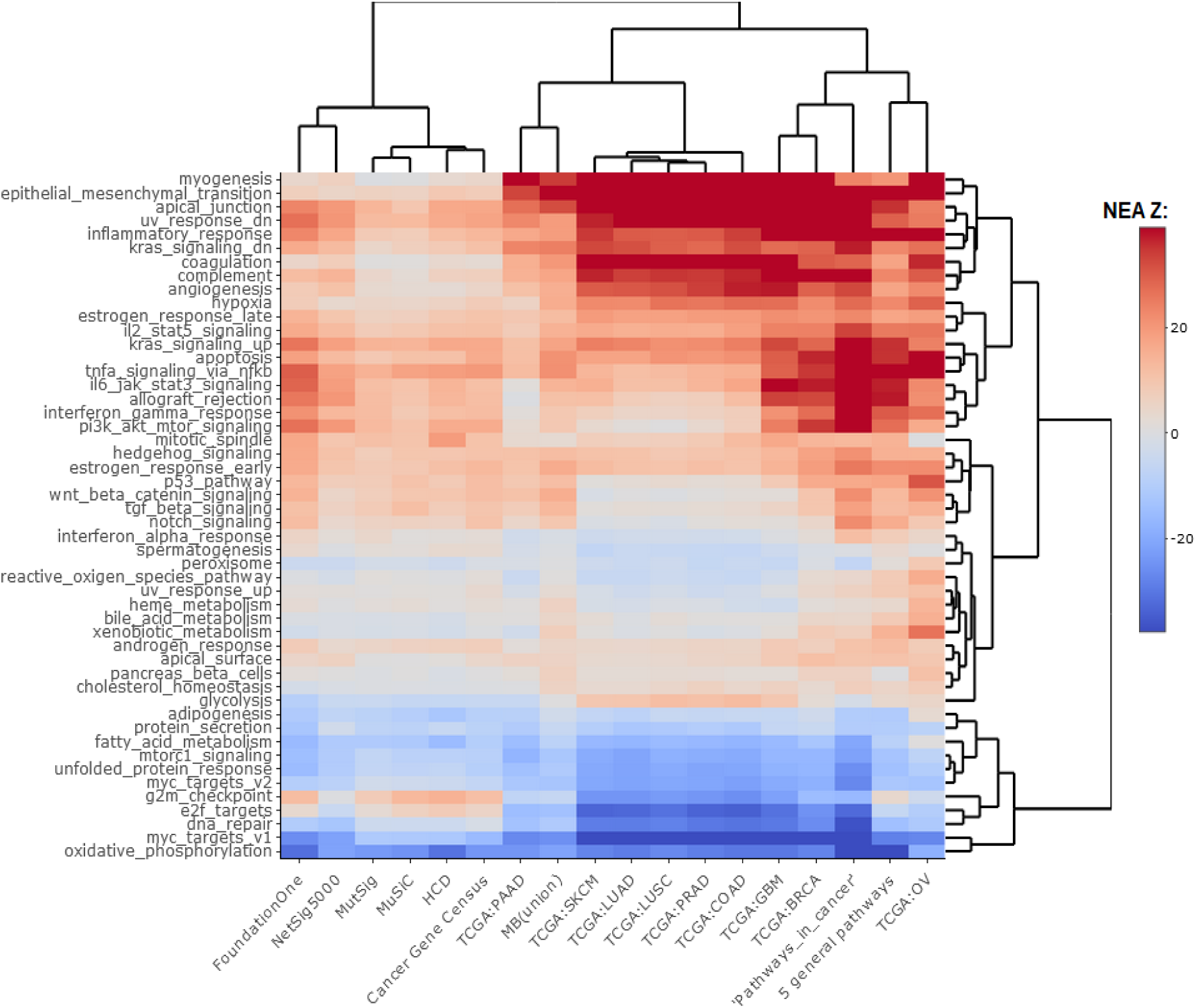
Agreement between NEAdriver and reference gene sets. A. The heatmap matrix elements represent overlap between the cohort-specific sets of predicted NEAdriver gene sets at q(M&P) < 0.05 and the gene sets from curated resources (lines 1…5) and alternative methods (lines 6…9). All the reference gene sets, except literature/KEGG site-specific pathways, had the same, “pan-cancer” membership independent of cancer site. B. Network enrichment of the cancer gene sets with regard to 50 hallmarks(41). NEAdriver sets defined at q(M&P)<0.05 are represented by 165 genes for each cohort, most frequent across its samples(n=165 was chosen for being half of the size of FoundationOne set).

Further, using NEA we positioned the different driver sets in the space of 50 hallmark gene sets(41) (Fig. 2B; interactive maps for q(M&P)<0.05 and q(M&P)<0.01: SupplFigure.drivers_vs_hallmarks.driverProbs.01 at https://rpubs.com/avalex99/748625 and SupplFigure.drivers_vs_hallmarks.driverProbs.05 at https://rpubs.com/avalex99/748627). It appeared that NEAdriver detected genes from a different hallmark subspace than the majority of the other computational methods. On the other hand, the NEAdriver predictions were clustered together with the curated sets KEGG05200 “Pathways in cancer” and the union of five “general” (not cohort-specific) cancer-relevant KEGG gene sets. Compared to the computational methods, the NEAdriver sets and the curated sets showed higher enrichment in EMT, angiogenesis, and suppressed KRAS signalling, glycolysis and hypoxia while were depleted in cell cycle, DNA replication/repair, peroxisome as well as MYC and mTOR signaling. For comparison, the original MGSs were more similar to the computational sets and differed from the curated sets (SupplFigure.drivers_vs_hallmarks.MGS at https://rpubs.com/avalex99/748620), which confirmed that the distinct NEAdriver pattern emerged from its specific features rather than reflected the initial mutation composition. We also noted that nearly all the alternative cohorts (except MutSig) abounded in genes with higher network degree, which were likely better known and more studied than the genes predicted with NEAdriver. Node degrees of the latter were closer to the network average, as illustrated by comparisons to random gene samples (Supplementary Fig. 2). Remarkably, the network-based method NetSig5000 also prioritized higher degree nodes.

### Estimation of discovery rates

The same reference gene sets were used for a more detailed evaluation of NEAdriver error rates. The best combination of true positive and true negative rates was found against the gold standard cohort-specific sets, which were either literature-based or available as KEGG pathways (Fig. 3A and 3B and upper left plots in Supplementary Figure 3), except BRCA cohort, where better results were found for NetSig5000 set which was derived from just this cohort(40).

**Figure 3.**
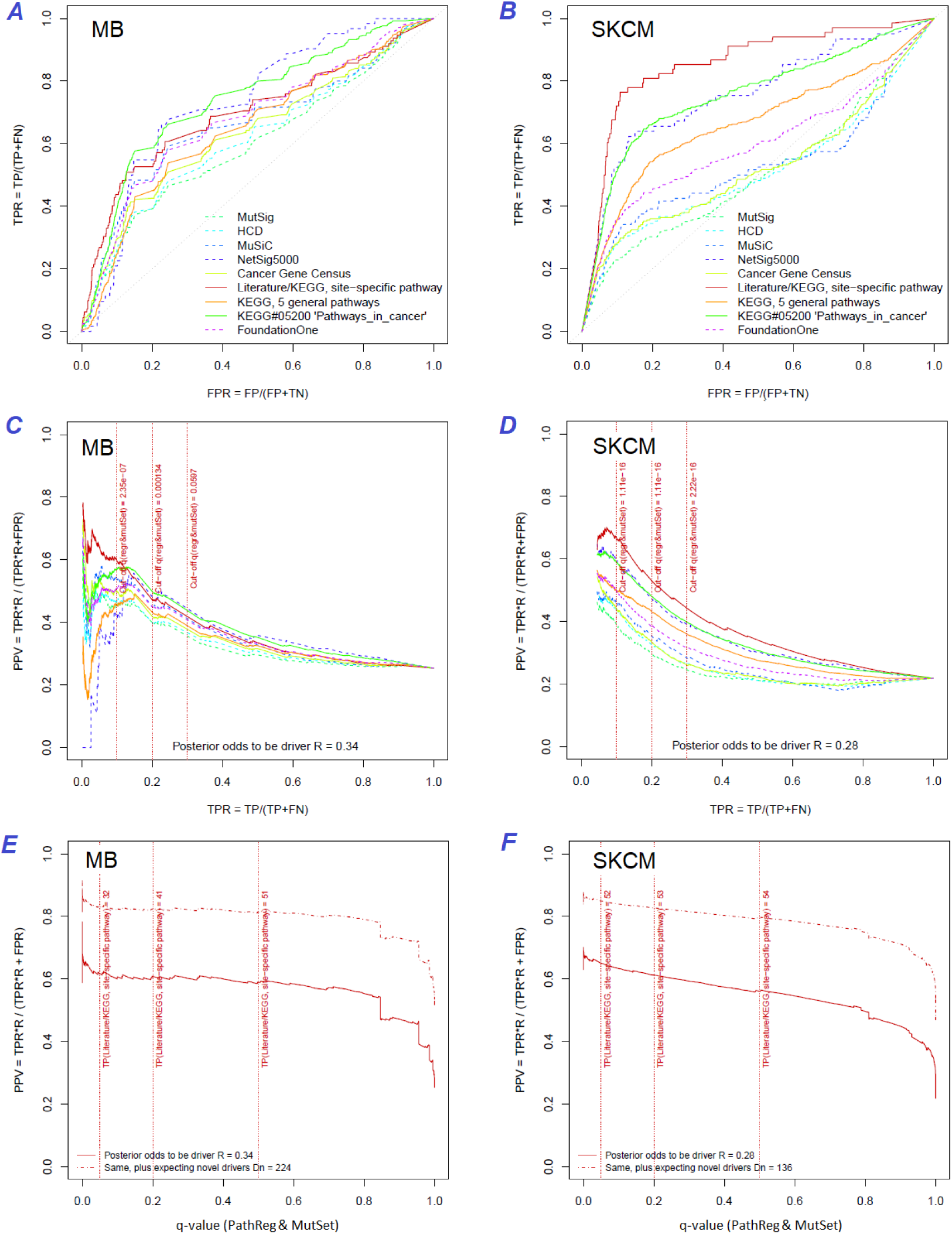
Performance of the new driver prediction evaluated on different benchmarks. Two cohorts with very low versus high passenger mutation load, medulloblastoma (MB: A,C,E) and skin cutaneous melanoma (SKCM: B,D,F), represent contrast conditions for computational driver discovery. The NEAdriver predictions were quantified by the cumulative statistic q(M&P)<0.05 and matched to nine reference sets. A and B: ROC curves in the space of true positive versus false positive rates in the classical definition of “precision”. C and D: Precision-recall curves where precision was calculated via inclusion of odds “driver/non-driver”. E and F: calibration of positive predictive value, PPV against false discovery rate (q ∼ 1 – PPV; solid lines) and modeling of PPV in presence of true, but yet unknown drivers (dot-dashed lines). The dotted vertical cutoff lines refer to the literature/KEGG, site−specific pathway sets. Cutoffs in C and D display q(M&P) values, whereas TP counts in E and F are numbers of unique “literature/KEGG” genes discovered under the respective q(M&P) threshold shown at X-axis.

Precision was first estimated using the common definition as fraction of true positives among all positives:

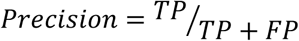

If applied to the full set of *n* genes, regardless of their mutation status in specific cancer genomes – which corresponded to GBA approach – then these estimates appeared very low and never exceeded 20% at TPR=10% (upper right plots in Supplementary Figure 3).

A more advanced estimate could be provided by using the hybrid positive predictive value (PPV) formula by John Ioannidis(42), where the frequentist terms – error rates of types I and II – were combined with odds (i.e. the ratio of actual drivers versus non-drivers among the mutated genes), which represented a Bayesian component:

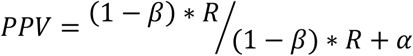

The odds could be estimated from the test results as

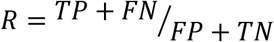

By expanding the Bayesian approach, the type I and type II errors would be, respectively:

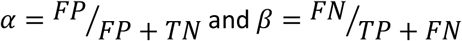

The value of FP+TN could be approximated as the total number of genes, since the number of drivers (TP and FN) should be negligible compared to that.

However, estimating the TP and FN via reference (gold standard) sets was more challenging, since the source publications and databases never claimed that their gene sets are truly complete. Thus, PPV estimates were particularly sensitive to biases in TP and FN and we therefore tried each of the nine sets. PPV ranged from 30 to 70% at TPR=10%, but even at TPR=100% almost never dropped below 20% (Fig. 3C and 3D and all cohorts in Supplementary Figure 3). Again, the best performance was achieved using the literature/KEGG sets (PPV=44…68% at TPR=10%).

Since this approach considered any genes not listed in each given set as false findings, the PPV estimates must have been excessively conservative. Therefore, we next investigated the potential of discovering novel drivers. As an example, applying this adjustment to the literature/KEGG sets under assumption that they were 50% complete increased PPV by 20…30% (dotted curves at Fig. 3E and 3F). Recalling that (1 – PPV) is essentially synonymous to false discovery rate (q-value) allowed us comparing error rate estimates from the two independent approaches: the gold-standard based PPV versus the continuous NEAdriver q(M&P). Although the relation was not linear over the range PPV=0…100%, at PPV=40…60% the estimates were remarkably close in each of the 10 cohorts (solid lines at Fig. 3E and 3F and Supplementary Figure 3).

We also compared the results to a number of previously suggested network-based methods that considered impact of somatic alterations on the transcriptome: DriverNet(43) and HotNet2(44) which implemented the cohort level approach as well as SCS(45), OncoIMPACT(46), and DawnRank(47) which worked at the individualized, single-patient level. The comparison also included naïve frequency-based estimates as provided by Guo and co-authors (Supplementary Figure 4). These publications presented short candidate driver lists combined over all samples. Agreement of the integrated ranks with NEAdriver confidence q(M&P) in four TCGA cohorts proved to be significant albeit rather weak (Spearman R=0.22…0.30). Further, we focused on lists of top 50 genes from each of the methods. In three cohorts (OV was the exception), a good agreement was found between the methods and NEAdriver (Supplementary Figure 5). Out of top 50 driver lists, between 9 and 41 genes received NEAdriver q(M&P)<0.05 (the overlaps were significant after Bonferroni-adjusted Fisher’s exact test p<0.001).

### Discovery rate vs. density

NEAdriver, designed to be independent from alteration frequency-, should be more sensitive to rare events compared to methods used such data. Indeed, the genes listed by Cancer Gene Census, FoundationOne, MutSig, HCD, and MuSiC had generally more mutation events per cohort than drivers predicted at *q(M&*P) <0.05 (Fig. 4). The same tendency was observed for the copy number alterations. The exceptions were SKCM cohort (which was not used as input by these projects), and few cohorts for MutSig (which implemented advanced normalization approaches). Sensitivity of NetSig5000 to rare mutations was comparable to our method – likely due to its network-based approach – but again mostly to genes with higher network degree (see details in Supplementary Figures 6 and 7). On the contrary, when mutation frequencies were normalized by coding sequence length, the differences between the methods vanished. This was not surprising, since shorter genes are less likely to mutate frequently.

**Figure 4.**
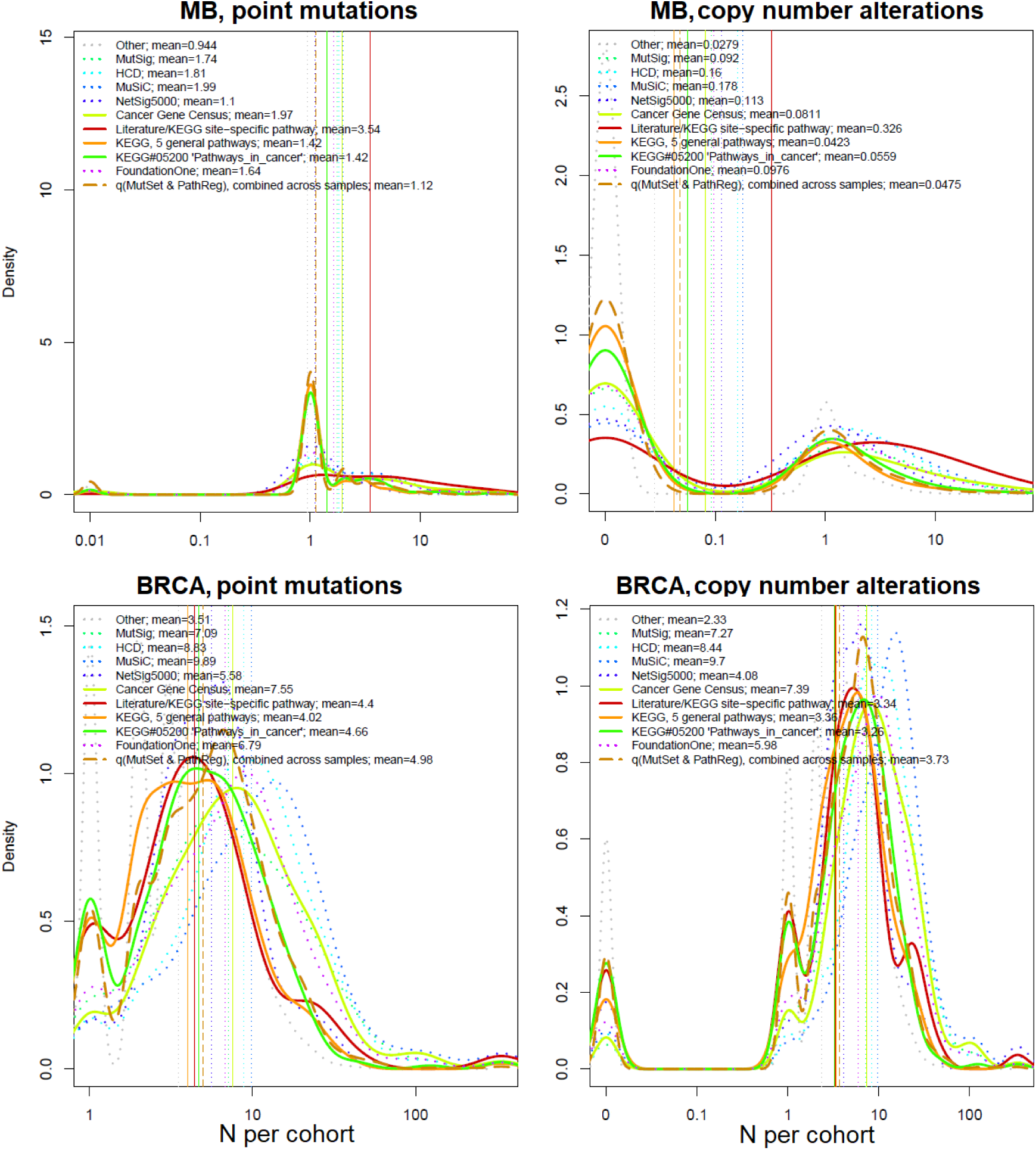
Comparative analysis of somatic alteration frequency among genes included in alternative reference sets. Density plots shape the distributions in each of the alternative sets, predictions by NEAdriver (q(M&P) < 0.05; dashed line), and genes not included in any of the above (“other”; dotted line). Vertical lines correspond to the mean values provided in the legend.

### Novel findings

How many known drivers there are in individual cancer genomes and by how much the new method could expand this space? An earlier computational analysis estimated the number of point driver mutations as two to six per genome(48). In our study – by counting any genes included in the nine alternative sets (N=1434) – the modes (most frequent count values) ranged across the cohorts between *M*=1…3 in MB (known to have very low somatic mutation load) to M=55 in SKCM (having typically thousands mutated genes per sample). For NEAdriver [q(M&P)<0.05], respective values per genome were lower, ranging between *M*=0…1 (MB) to *M*=50 (SKCM) (Fig. 5A). Overlaps between these two approaches were rather modest (*M*=0…8). In other words, the driver candidates identified by NEAdriver were mostly novel. The overlaps between “alternative sets” and NEAdriver [q(M&P)<0.01] are also presented for individual cancer genomes (Fig. 5B). The Jaccard coefficient values, with exceptions of MB and GBM, rarely exceeded 0.3, which confirmed that NEAdriver identified mostly novel genes.

**Figure 5.**
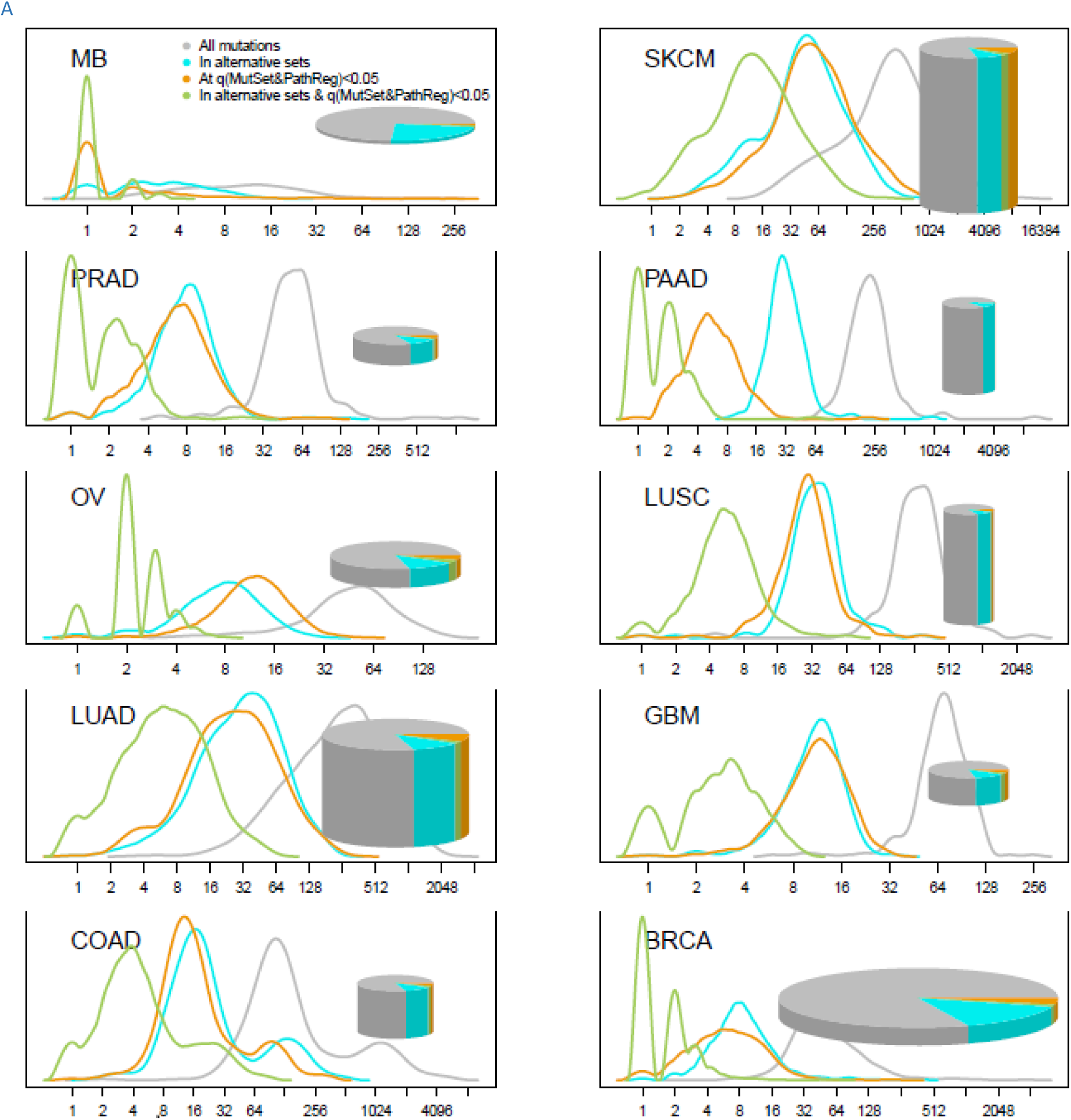
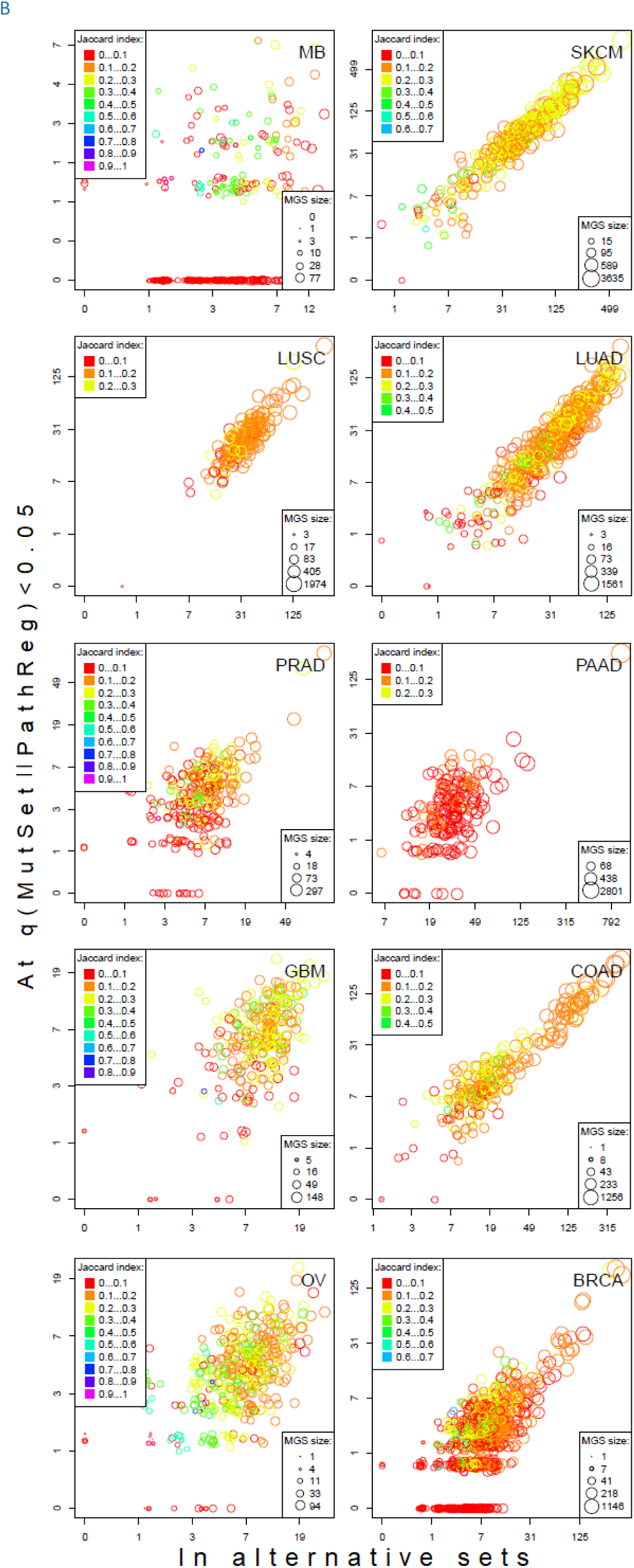
Distribution of somatic mutations versus drivers across genomic samples. A. Relative density plots of mutations and declared drivers. Pie charts summarize counts per genomic sample in each of the ten cohorts (height: average number of reported mutations per sample; width: number of samples in the cohort). B. Overlap between the predictions by MutSet|PathReg and the merge of alternative gene sets (1434 genes in total) measured with Jaccard index values (sets’ intersection divided with sets’ union), denoted by color. The MGS sizes (regardless of driver status) are denoted by marker size. Gaussian noise was added to marker coordinates for better readability.

Recalling that respective PPV estimates reached 50% and exceeded 75% when allowing for novel drivers, the predictions appeared fairly confident.

### Clustering patient driver sets in pathway space revealed association with survival

One of the goals of tumour molecular profiling is to discover cancer subtypes which would be informative of disease outcome or clinically meaningful otherwise. The driver genes identified with our method were mostly rare and therefore not suitable as stand-alone subtype markers. However, using NEA we could generate “DGS vs. FGS” scores which summarized signals from various disparate events and thus available for every patient. We explored if DGS profiles in the FGS space could partition the cohorts by differential survival.

Indeed, the DGSs [q(M&P)< 0.05] were often informative on patient survival. We tested three different clustering techniques and found that in many cases DGS scores differentiated cohorts by survival: 7…14.8% of all tests yielded significant Cox proportional hazard models (Benjamini-Hochberg FDR < 0.25). Furthermore, in up to 21.1% of all tested cases the significant partitions were recapitulated on test sets (while FDR estimates from Cox models were below 0.25) (see examples in Fig. 6 and full details in Suppl. Fig. 8). For comparison, splitting in the same framework by high vs. low tumor stage did not differentiate patients by survival (not shown).

**Figure 6.**
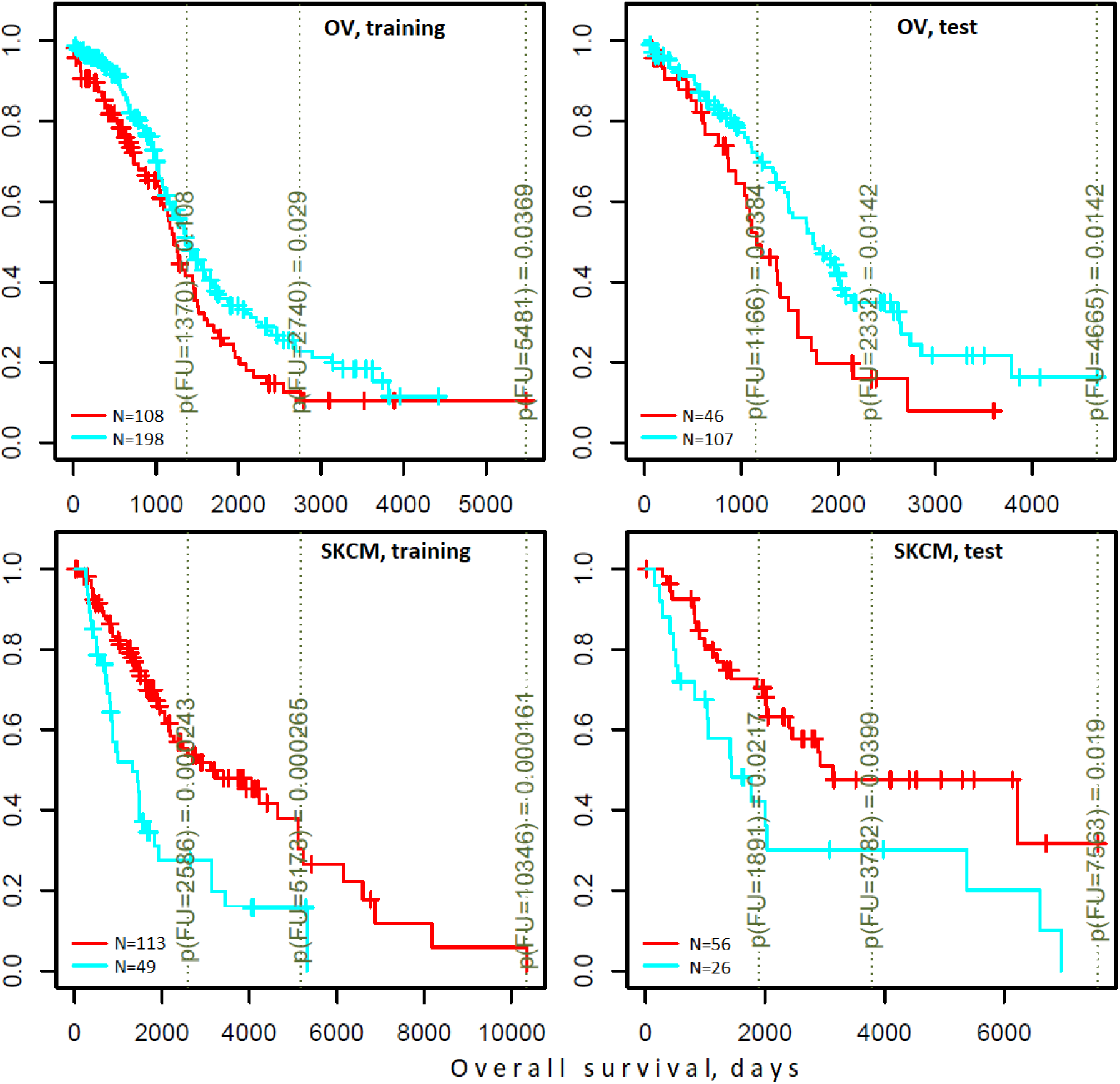
Differential survival of patients stratified in pathway space created by network enrichment analysis of driver gene sets. Vertical captions (brown) convey Cox proportional hazard p-values for three follow-up intervals.

### NEA scores based on either drivers or gene expression point to same pathways associated with survival

Finally, we checked if association of specific FGS scores with survival could be traced at the level of mRNA transcription. To this end, we derived lists of 100 patient-specific genes with expression most deviating from the cohort mean (GE.AGS) and looked if their NEA scores for the same FGS would also be associated with survival. By testing the 10 cohorts, 2 survival types, 3 clustering methods, and the 1659 FGSs, we identified 31 cases where the association with survival was observed for both DGS-FGS and GE.AGS-FGS scores. The discovery of this many associations was significant in a random permutation test requiring Bonferroni-adjusted p-value < 0.01 while permutation test-based p-value < 0.0001 (Suppl. Fig. 9). Remarkably, in most of the cases DGS and GE.AGS demonstrated opposite relations with survival: better outcome was associated with high scores of the former while lower scores of the latter, or vice versa (Supplementary Figure 10).

For example, MB and LUAD cohorts were differentiated by survival using NEA score profiles for two pathways (Fig. 7). The cohort patients were represented first by DGSs (left) and then by GE.AGSs (right). The NEA scores reflected connectivity between pathway genes and patient genes (either DGS or GE.AGS). A higher NEA score would indicate that relatively many patient-specific genes were linked to the given pathway. MB cells are known to sometimes produce granulocyte colony-stimulating factor (49), which can affect influx of granulocytes(50) and disease prognosis(51). With regard to “Biocarta granulocytes pathway”, the MB patients where stratified so that higher DGS scores indicated poorer survival, whereas higher GE.AGS scores were associated with better survival. Subnetwork patterns for two patients exemplify this analysis (Fig. 7A). ALK fusion events are a well-established target for non-small cell lung cancer therapy(52). While none of the patients were treated with an ALK inhibitor in LUAD cohort, “Biocarta ALK1 pathway” scores for both DGS and GE.AGS were informative on overall survival within 6-year follow-up interval. Again, relations to survival were opposite for DGS versus GE.AGS scores.

**Figure 7.**
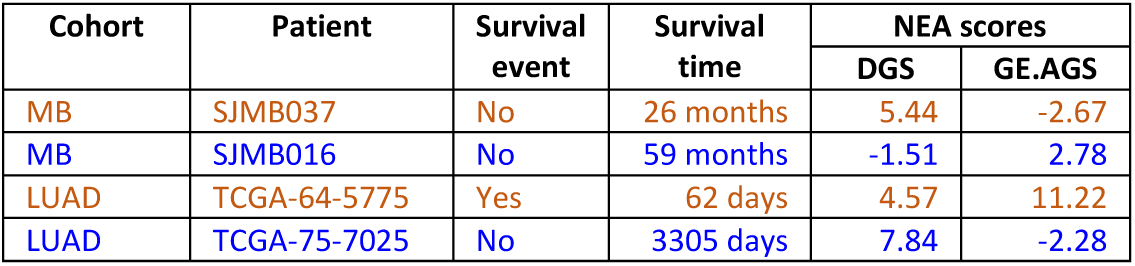
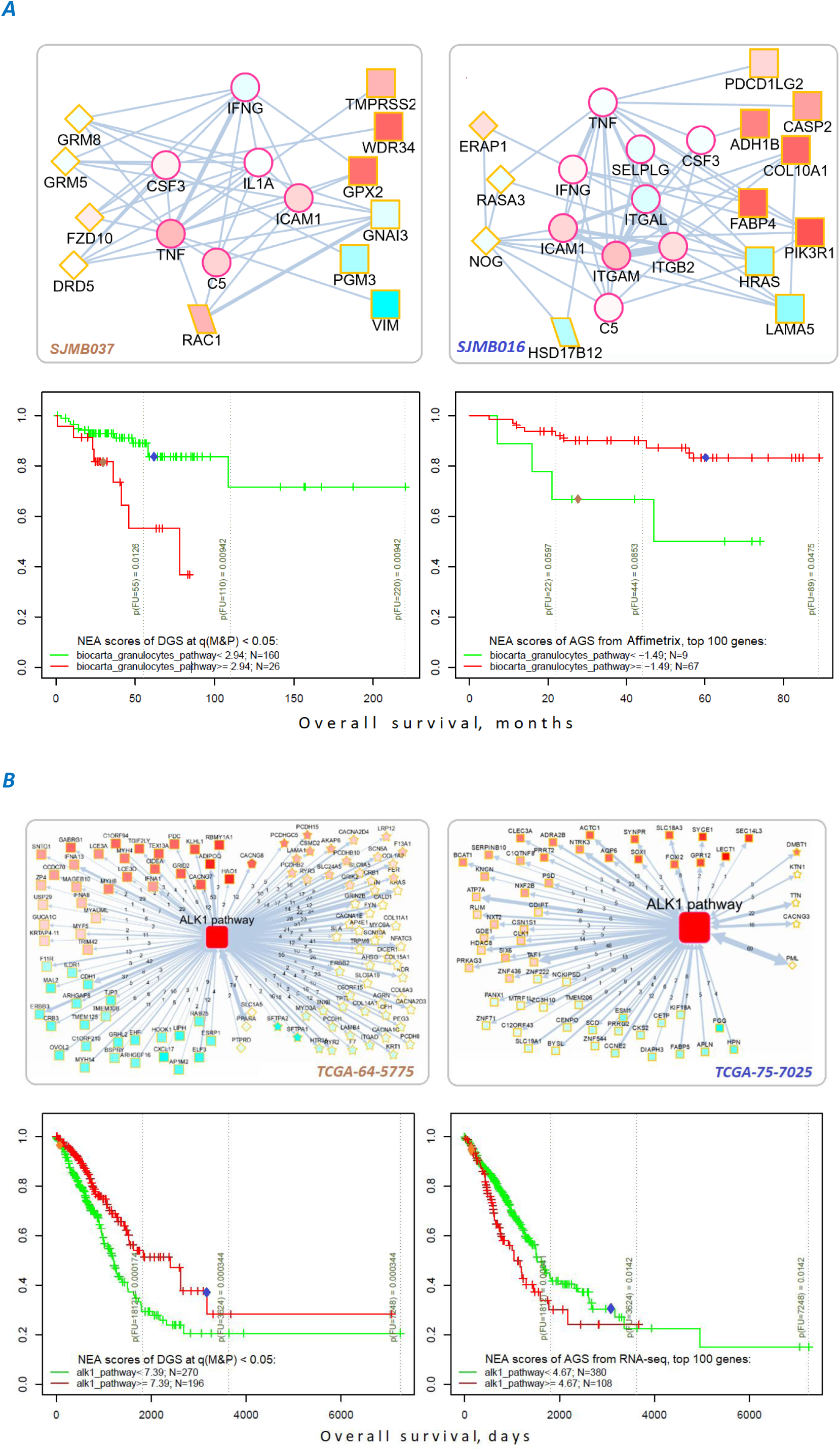
Network enrichment and survival analyses of patient specific lists of drivers and differentially expressed genes. **A. Example from MB cohort**. **B. Example from LUAD cohort**. Yellow borders: patient-specific gene sets including 1) driver alterations (*q(M&*P)<0.05): either point mutations (stars) or copy number changes (diamonds) 2) genes with mRNA expression most deviating compared to the rest of the cohort (rectangles) 3) both categories 1 and 2 (rhomboids) Magenta borders: pathway genes (circles). Each gene is colored by expression in the given patient sample compared to the cohort mean. Note that pathway genes usually did not manifest genomic or strong expression changes. **In figure B**, the edges combine individual network links between genes. Links within pathway not shown. **Clinical and NEA data for the patients:**

## Discussion

The need in novel approaches for driver identification is urgent: while every cancer genome is expected to possess driver mutations, many cases would lack any alterations in known cancer genes – which is counterintuitive and undermines the ground for targeted therapy. Due to the high mutational heterogeneity of cancer samples, frequency-based methods have reached their limits of statistical power to detect novel cancer drivers. Network analysis – an already popular method to identify cancer genes using functional context – in our implementation was relatively less biased toward network hubs and thus more sensitive to novel driver genes. The previous guilt-by-association analyses(30,53–55) predicted gene function based on functional connections to known functional categories. Confidence of such predictions alone, e.g. in absence of experimental data was very low due to rare occurrence of actual mutations and thus lower true discovery rate, as shown by John Ioannidis(42). NEAdriver was more focused due to considering concrete molecular phenotypes (*de facto* alterations in individual genomes) and combining relevant evidence from two network analysis channels.

Since capabilities of both `MutSet` and `PathReg` could only be implemented on driver constellations of sufficient size, it was important to test NEAdriver on cancer genomes with different mutation loads. This feature varied from a few affected genes in MB to thousands in lung and skin cancers. Despite the variability of this and other biological parameters, both statistical performance and candidate drivers’ functional profiles proved to be rather close across the ten cohorts. As an example, three pathways were systematically included in the models: hsa04020:Calcium_signaling_pathway (CS), hsa05412:Arrhythmogenic_right_ventricular_cardiomyopathy_(ARVC), and hsa05414:Dilated_cardiomyopathy (DCM). Although the role of CS in cancer was rarely considered central, cell migration and adhesion do involve modulation of cell motility and shape where ion channels and pumps play major roles, so that CS genes are known for both downregulation and functional implication in cancers(56),(57),(58). ARVC and DCM are functionally close to CS, although have little overlap in member genes. At the first glance, the major factor behind MGS-CS interrelations might be the frequently mutating titin TTN. Its involvement in cardiopathies and cancer has been long argued because of its extremely long coding sequence, thus likely prone to spurious alterations (∼50% of LUAD samples). However, there were many individually rare mutations, which together revealed emergent network patterns between MGSs and either CS, or ARVC, or DCM: cadherins, laminins, integrins, metalloproteases, nitric oxide synthases, ryanodine receptors, adenylate cyclases, subunits of protein kinase A etc. They contributed to the NEA scores with multiple network links so that e.g. the median edge counts between genes of MGSs and of the pathways were 22…45 in MB and 497…890 in LUAD.

The cohort- and sample-specific NEAdriver predictions agreed well with nearly all tested alternative sets (Fig. 2A). The best agreement was found for cancer site specific gold standard sets while less so for pan-cancer sets. The computational, mostly frequency based sets performed worse than curated sets (Fig. 3A,B). In the functional space, NEAdriver predictions were positioned differently compared to computational and database sets, but close to curated cancer pathway sets (Fig. 2B), which confirmed both novelty and relevance of NEAdriver findings. The latter also differed from most of the alternative sets in having much less bias in regard of network node degrees and gene length.

A realistic estimate of the true discovery rate for NEAdriver predictions was obtained by accounting for putative drivers not included of the gold standard sets, so that PPV could be as high as 78%…88% at the q(M&P)=0.05, which was validated by the two alternative way of PPV calculation (Fig. 3E,F). The efficiency of NEAdriver was confirmed by the ability of DGS to stratify patients by survival (Fig. 6) and the striking tendency of same pathways being associated with survival via both mutation- and expression-based patient scores (Fig. 7).

Obviously, NEAdriver alone would miss events detectable by other methods, e.g. when certain “stand-alone drivers” impose strong effects on their own, without apparent interaction with other genes. This creates an incentive for creating combined methodology and a toolbox in the future. But already now our analysis identified hundreds of somatic gene alterations that had not been deemed functional in previous research. After evaluation performed in a number of ways, the body of predictions appears confident, providing a set of provable research hypotheses and suggesting new strategies for cancer prognosis and individualized treatment.

## Materials and methods

### Medulloblastoma meta-cohort

We collected data from publications presenting large-scale datasets (59),(60),(6),(61) and two public datasets available online (PBCA-DE and PEME-CA). We retrieved available exome sequencing profiles as well as copy number alterations, gene expression, and clinical data. We translated gene identifiers into gene symbols according to ENSEMBL annotations v.93 and then made sure all the gene symbols are found in the network and are up to date according to GeneCards(62) annotations.

For consistency with the publication datasets, we excluded the following types of mutations from PBCA-DE and PEME-CA sets: intron variant, upstream gene variant, 3_prime_UTR_variant, 5_prime_UTR_variant, intergenic region, downstream gene variant, synonymous variant, and splice region variant. For a few patient IDs that were found in more than one dataset, their mutation profiles were merged (if different).

Overall survival data was collected from the published datasets. A few patients with discrepant data (for instance, ICGC_MB193 was 2.3 years old according to Northcott dataset, but 70 years old according to PBCA-DE dataset) were excluded. For 18 samples with different follow-up we accepted the newest survival time values from Northcott dataset.

Data from all the datasets were combined into one cohort dubbed MB(union), so that 541 patients were covered with both clinical and exome sequencing data.

### TCGA cohorts

The TCGA^63^ data were obtained via https://portal.gdc.cancer.gov/.

### Network Enrichment Analysis

Network enrichment between two gene sets of interest *S*_*a*_ and *S*_*b*_ is estimated by comparing the actual number of network edges 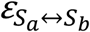 that connect nodes of *S*_*a*_ with nodes of *S*_*b*_ in the real, biological network *G*_*B*_=(*E,V*) with a number expected by chance 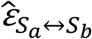 in a random network *G*_*R*_=(*E,V*) where particular node degrees *k* of genes 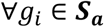; 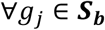 *g*_*i*_ ≢ *g*_*j*_ equal to those of the actual network (which implicitly assumes that the whole degree sequences of *G*_*B*_ and *G*_*R*_ are identical, too). In an earlier work (37), series of randomized instances of *G*_*R*_ were created using an algorithm of explicit edge permutation (63) and used for estimating expected variance of *ε*. Later, it was demonstrated (38) that 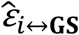 can be calculated analytically in a fast and unbiased manner:

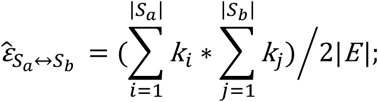

Then the difference between the actual and expected edge counts

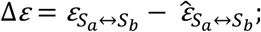

is used to estimate significance of the relation *S*_*a*_↔*S*_*b*_ with a *χ*2 statistic:

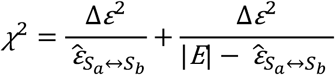

The *χ*2 does not follow Gaussian distribution, but it can be conveniently converted to Z-scores and then used safely for downstream calculations in e.g. linear models.

In the simplest NEA case one of the sets is a single gene *i*:

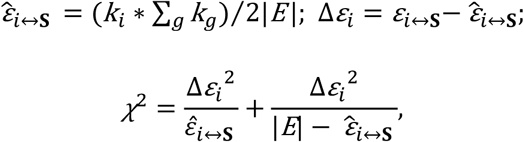

which simplified calculation and – within this work – enabled estimation of network enrichment for a mutated gene against the (rest of) mutations in the same cancer genome, called mutated gene set (MGS) in MutSet method or functional gene set (FGS, or simply pathways) in case of PathReg

### Network

For NEA we merged network of top 1 million edges, ranked by confidence (i.e. Final Bayesian Score) from FunCoup 3 (Schmitt et al., 2014) and all edges of Pathway Commons 9(64,65). We made sure that all genes reported as altered in at least one of the ten cancer cohorts had up-to-date gene symbols in network. That resulted in a network of 19,035 nodes (unique gene symbols) connected with 1,731,648 unique edges.

### Mutation gene sets and driver gene sets

We defined mutated gene sets (MGS) as lists of all genes of a given tumour sample reported with somatic mutations (SM) in the MAF files. MGSs were used as whole sets in driver evaluation of MutSet channel. MGSs ***did not include*** copy number altered (CNA) genes.

For TCGA, the analysis included all mutations reported in the MAF files, regardless of predicted functional impact. Indeed, although synonymous and intronic mutations were often disregarded in cancer research, their involvement in carcinogenesis seems likely and has been recently demonstrated (66). In our datasets frequency of silent mutations was somewhat lower than of non-synonymous ones, but still significant so that many frequently mutated genes showed elevated rates in both categories (Supplementary Figure 11). Therefore, each altered gene *i* from a given sample, either SM or CNA, was evaluated against the MGS and received a MutSet q-value. Significantly altered genes with were included in final driver gene sets DGS_0.05_ and DGS_0.01_ under conditions *q(M&*P) < 0.05 and *q(M&*P) < 0.01, respectively.

### Functional gene sets

For the PathReg predictor, we used 318 KEGG pathways from version as of 16 August 2018. Considering the importance of SHH and WNT pathways in e.g. medulloblastoma, alongside with respective KEGG pathways we included these two also in Biocarta (67) versions (the versions were very different in size and length). We updated gene symbols in the sets in the same way as described for the mutations above.

For the survival analysis, the FGS collection consisted of 1659 entries from BioCarta (67), KEGG (68), Reactome (69), WikiPathways (70), MetaCyc (71), and MSigDB hallmarks (41).

### Altered gene sets (transcriptomics)

The gene expression data was used from the available cohort data sets:

- Affymetrix for MB (61);
- Agilent for OV and GBM;
- IlluminaHiSeq_RNASeqV2 for the rest of TCGA cohorts.

The AGS were compiled as sample-specific lists of top *N* genes (*N*=[50,100,200]) with normalized mRNA expression most different from the respective cohort mean using function `samples2ags(…, method = “topnorm”)` from R package `NEArender` (38).

### NEAdriver: algorithm

The driver discovery algorithm combined results from two NEA-based channels, `MutSet` and `PathReg`.

#### MutSet channel

The `MutSet` values quantified network enrichment between each gene *m* having a somatic point mutation in genome *j* and the set MGS of all other point mutations in the same genome (*m* ∈*MGS*_*j*_). They were calculated as NEA scores 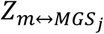 so that within a cohort the same gene might receive multiple, sample-specific `MutSet` values.

#### PathReg channel

As the independent variable for training the `PathReg` predictor, we employed vectors `anchor.summary`. Specific NEA scores were calculated for every gene *i* present in the network (*N*=19035) versus every MGS in the given cancer cohort *c*. The `anchor.summary` values *µ*_*ic*_ were then obtained by summing up over all *N*_*c*_ available samples, regardless of mutation status of *i* in genome *j*:

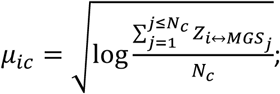

Since the score 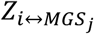 is derived from the network patterns of mutated genes across the cohort and does not depend on the mutation profile of *i* itself, the *µ*_*ic*_ value would reflect a general propensity of *i* to interact with constellations of putative cancer genes. The transformations via χ2 ⟶ Z, log, and square root were imposed in order to render distributions closer to Gaussian.

The *µ*_*ic*_ profiles were rather scarce due to rare occurrence in MGS of true drivers that would interact with a given gene *i*. We thus further improved the gene specific values via modelling *µ*_*ic*_ with pathway NEA scores *Z*_*i*↔*FGS*_. These were calculated for 320 FGS versus each of the *N* network genes and then used as a matrix of dependent variables **Φ** in `PathReg` training.

Then sparse regression models were created using function `cv.glmnet` from R package `glmnet` (72).The chosen package implements elastic net models for solving the problem:

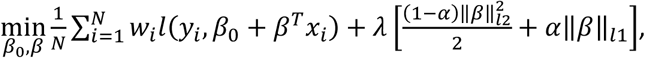

where α is a mixing parameter for balance between lasso and ridge regression (whereby α=0 and α=1 would lead to plain ridge and lasso regressions, respectively). In our case (α=1), glmnet solved just the lasso problem:

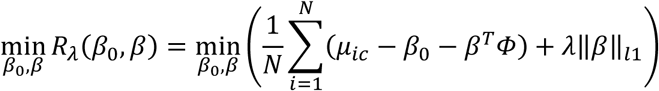

Parameter λ determines complexity of the multivariate regression model, i.e. what subset of initially submitted variables of **Φ** should receive non-zero coefficients. Under 3-fold cross-validation, function `cv.glmnet` tested a series of λ values while controlling the cross-validation mean squared error (CVM). The cohort-specific optimum *λ*_*с*_ was found as a trade-off between model precision and complexity using Bayesian information criterion (BIC), which was deemed preferable(73) over Akaike information criterion in the context of favourable dimensionality (*n*_*m*_ ≫ *p*;*p* = 320). The optimal *λ*_*с*_ was set at the number of FGS variables with non-zero coefficients *k* as the smallest possible within 2 standard errors of BIC from the lowest BIC value:

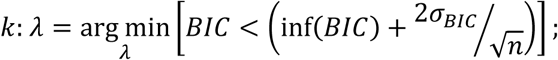

At these *λ*_*с*_, the values of CVM and residual sums of squares were close to their respective minima, too. The obtained regression models explained *µ*_*ic*_ with linear combinations of FGS NEA scores *f* = *Z*_*i*↔*FGS*_:

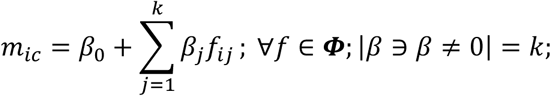

After this training and model selection step, the retained test subsets were used to check how the original values *µ*_*ic*_ correlate with the predicted values *m*_*ic*_ (Supplementary Fig. 1).

The distribution of *m*_*ic*_ values was non-parametric but close to Gaussian. Therefore, respective p-values were modelled via a normal distribution where mean and standard deviation were estimated as median and 84.2^th^ percentile of the empirical distribution, respectively:

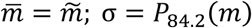

(in the Gaussian distribution 84.2% of values are within 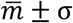).

#### Integration of channels

Both `MutSet` and `PathReg` produced output in the form of *q-*values, equivalent to false discovery rate which conveys the probability of a given driver prediction to be false. These values were integrated into the final value as a product *q(M&*P)=*q*_MutSet_\**q*_PathReg_, which presented the probability that neither channel have produced true predictions. Therefore, 1 – *q(M&*P) was the probability of either channel to be true and we convened to trust a driver prediction if *q(M&*P) < *c* (*c*=[0.01, 0.05]).

### Gold standard and alternative driver sets

#### Literature based sets

As gold standard for **MB**, we compiled a list of unique 516 gene symbols, of which 12 were found in OMIM database, 140 in Disease Ontology, and 399 in MB-related publications found in PubMed:

We included all altered genes mentioned in these publications. Mechanisms of alteration included changed methylation, gene copy number, and point mutations.

For the TCGA cohorts, we employed the dedicated KEGG pathways:

**BRCA** <- hsa05224:Breast_cancer;

**GBM** <- hsa05214:Glioma;

**LUAD** <- hsa05223:Non-small_cell_lung_cancer;

**LUSC** <- hsa05223:Non-small_cell_lung_cancer;

**SKCM** <- hsa05218:Melanoma;

**PRAD** <- hsa05215:Prostate_cancer;

**PAAD** <- hsa05212:Pancreatic_cancer;

**COAD** <- hsa05210:Colorectal_cancer;

**BLCA** <- hsa05219:Bladder_cancer;

**OV** <- hsa05213:Endometrial_cancer, since origins of these two are intertwined(74).

## Sets from computational analyses

**MuSiC, N=127**(48) identified drivers based on relative point mutation frequency, assisted with expression analysis and database annotations in 12 TCGA cancers (of which six overlapped with our analysis). We used the “PanCancer” list.

**MutSig, N=260**(14) performed a comprehensive point mutation frequency analysis using exome sequencing data from 21 cancer cohorts (of which nine overlapped with our study), while accounting for mutation burden, clustering, and functional impact.

**HCD, N=291**(21) discovered drivers in 12 TCGA cohorts (of which six overlapped with our analysis)with a combination of four algorithms that prioritized mutated genes based on mutation rate, functional impact, positional “hotspots”, and specific enrichment in phosphorylation sites.

**NetSig5000, N=62**(40) gathered evidence for potential driverness of each gene via functional coupling to frequently mutated genes in the global network. The genes’ own mutation frequencies were incorporated into NetSig scores at a separate step.

## Pan-cancer database sets

**Cancer Gene Census N=270**(75) was downloaded on 7^th^ of February, 2019. The genes from the ten cohorts were merged in a single list.

**KEGG#05200:Pathways_in_cancer, N=395**(68) was a “pan-cancer” version including a curated selection of organ-specific cancer pathway gene lists.

**Five “general” cancer pathways, N=457**(68) was created as a union of the following cancer related KEGG pathways:

hsa05202:Transcriptional_misregulation_in_cancer;

hsa05203:Viral_carcinogenesis;

hsa05204:Chemical_carcinogenesis;

hsa05205:Proteoglycans_in_cancer;

hsa05206:MicroRNAs_in_cancer.

**FoundationOne, N=330** (https://www.foundationmedicine.com/resources) is the targeted sequencing panel used for cancer diagnostics, created as a merge of “general” and “rearrangements” sections.

## Sets from individualized analyses

**OncoIMPACT (N=162…695 per cohort)**(46) used an expression-driven approach to validate driver roles of point mutations in five TCGA cohorts (of which four overlapped with our analysis). The paper reported cohort-specific driver ranks rather than individual, sample-level estimates. The genes that received a rank were used in comparisons with NEAdriver results.

**SCS**(45) reported driver role evaluation in individual samples. They reported only top 50 genes after global, cohort-specific ranking. In parallel, the tables contained **top 50 genes** from **OncoIMPACT**(46), **DriverNet**(43), **DawnRank**(47), and **HotNet2**(76)(44) methods, which we also imported and used in our comparison.

## Normalization of mutation frequencies

The number of samples where each given gene was mutated was normalized by dividing it with its CDS length.

## Acknowledgements

The authors are grateful to Vetenskapsr**å**det for provided funding. The analysis used data generated by the TCGA Research Network: https://www.cancer.gov/tcga.

## Author Contributions

AA conceived the study and the algorithms. IP performed the data collection and curation. Both authors performed the analysis and wrote the manuscript.

